# Elevated environmental CO_2_ stimulates photosynthesis and induces differential oxidative responses in two Caribbean reef-building coral species

**DOI:** 10.64898/2026.10.16.759725

**Authors:** M. Tresguerres, S.E. Sabatini, L.B. Linsmayer, C.T. Galindo-Martínez, A.B. Thies, T.J. Hamilton, D. I. Kline

## Abstract

Reef-building corals maintain intracellular photosymbiosis with dinoflagellate algae, allowing them to use photosynthetic products as metabolic fuel. Carbon dioxide (CO_2_) is a key substrate for photosynthesis, but its rapid increase in the global ocean due to anthropogenic emissions (known as “ocean acidification”) may disrupt coral symbiosis. We examined the effects of elevated CO_2_ on photosynthetic O_2_ production, oxidative damage, and antioxidant systems in the Caribbean reef-building corals *Porites astreoides* and *Orbicella franksi*. Coral colonies collected at 3m depth in Bocas del Toro (Panama) were exposed to control (∼390 μatm CO_2_, pH ∼8.00) or elevated CO_2_ (∼1,060 μatm CO_2_, pH ∼7.70) conditions in outdoor flow-through aquaria for 10 days. Flowing seawater had higher O_2_ concentrations in elevated CO_2_ tanks, with the largest differences at higher irradiance. Microsensor measurements confirmed that acute and chronic exposure to elevated CO_2_ significantly stimulated photosynthetic O_2_ production in both species. Despite increased O_2_ production. neither species showed significant changes in lipid peroxidation, reduced glutathione, or superoxide dismutase, catalase, and glutathione-S-transferase activities. Protein oxidation was unchanged in *O. franksi* but significantly decreased in *P. astreoides*. These results are consistent with partial relief of carbon limitation in *P. astreoides* symbionts under elevated CO_2_, potentially improving photosynthetic efficiency and decreasing reactive oxygen species production. These findings support the resilience of *P. astreoides* to OA and suggest that elevated CO_2_ may increase daytime O_2_ levels on future reefs, with implications for the aerobic metabolism of reef organisms.

## Introduction

Tropical coral reefs are biodiversity hotspots that provide myriad ecosystem and economic services. The reef framework is largely sustained by cnidarian animals that establish an endophotosymbiotic association with dinoflagellate Symbiodiniaceae algae. These algae produce sugars and other nutritional compounds that supply the majority of carbohydrate needs for the cnidarian hosts and are essential for their thriving in oligotrophic tropical seas. However, anthropogenic activities that result in local and global stressors including ocean acidification and deoxygenation may threaten the health of the cnidarian-algal symbiotic association and, by extension, that of coral reefs (Nelson and Altieri, 2019; Pandolfi et al., 2011; Pezner et al., 2023). Ocean acidification is caused by increased absorption of atmospheric CO_2_ into the ocean leading to decreases in average surface seawater pH and [CO_3_^2-^] and an increase in [HCO_3_^-^] (Feely et al., 2004). The initial research in the early 2000s predicted catastrophic consequences of ocean acidification on carbonate biomineralization (Albright et al., 2016; Andersson and Gledhill, 2013) and studies over the subsequent two decades have generally confirmed increased dissolution rates of exposed reef carbonate and species-specific effects on biomineralization (Leung et al., 2022). The effects of ocean acidification on coral photosynthesis are similarly variable but it is clear that, at least in some species, elevated CO_2_ may enhance photosynthetic activity (Armstrong et al., 2025; Biscéré et al., 2019; Castillo et al., 2014; Langdon and Atkinson, 2005; Martins et al., 2024; Mason et al., 2020), possibly by relieving carbon limitation. However, other studies have reported that elevated CO_2_ may overwhelm cnidarian antioxidant responses, leading to increased oxidative damage (Jiang et al., 2019; Luz et al., 2018; Siddiqui and Bielmyer- Fraser, 2015; Soriano-Santiago et al., 2013). This contrasts with the hypothesis that carbon limitation itself promotes oxidative (Wooldridge, 2010; Wooldridge, 2013), in which case relief of carbon limitation under elevated CO_2_ could instead decrease oxidative damage. However, comparing results from different studies and inferring generalized responses is a challenging task due to the heterogeneity of experimental coral species, time in captivity, seawater quality, light and flow levels, feeding regime, duration of the exposures, and other details of the techniques utilized in the various studies.

We conducted an integrative study that quantified the effects of elevated CO_2_ on oxidative stress and antioxidant responses of two Caribbean coral species, *Porites astreoides* and *Orbicella franksi.* These species were selected due to their different ecological characteristics and reported physiological responses to environmental change. *P. astreoides* is considered a climate change “winner,” thriving in a wide range of habitats and often maintaining photosynthetic performance and oxidative balance under elevated CO, (Bove et al., 2019; Enochs et al., 2016; Lima et al., 2022). *O. franksi* is a historically important reef-building Caribbean coral whose physiological responses to environmental stress have also been extensively studied (Linsmayer et al., 2024; Silbiger et al., 2019). While previous studies often examined these species in different locations, times, environmental contexts, and artificial aquarium settings, here we directly compared their physiological responses by collecting both species from ∼3 m depth at Punta Caracol reef (Bocas del Toro, Panama) and maintaining them in an outdoor aquarium system supplied with flowing seawater from an adjacent reef site. This approach allowed us to study species-specific responses to elevated CO_2_ in an environmentally relevant setting that matched natural temperature and light dynamics.

During routine monitoring of the experimental aquarium, we noticed that the coral-holding tanks from the elevated CO_2_ treatment tended to display higher dissolved O_2_ levels than the coral- holding tanks from the control treatment, a trend that was especially pronounced on sunnier days. This observation prompted us to explore coral photosynthetic O_2_ production rates in control and elevated CO_2_ conditions using O_2_ microelectrode measurements on the diffusive boundary layer (DBL).

Our results showed enhanced photosynthetic O_2_ production in both *P. astreoides* and *O. franksi* during a 10-day exposure to elevated CO_2_ and suggest that this response has the potential to increase O_2_ levels in the bulk seawater overlying a reef during the daytime. And while antioxidant defenses, coloration, and lipid peroxidation levels remained unchanged in both species, *P. astreoides* experienced reduced levels of protein oxidation in the elevated CO_2_ treatment. In addition to helping explain the known resilience of *P. astreoides* to ocean acidification, these results have interesting implications on the aerobic metabolism of corals and coral-associated organisms in future high CO_2_ oceans.

## Methods

### Specimens

During October 10-15, 2016, coral colonies (∼10 cm diameter) of *P. astreoides* and *O. franksi* were collected while SCUBA diving at 3 m depth in the Punta Caracol Reef, Bahia de Almirante, Bocas Del Toro, Panama (9.363°N, 82.282°W). Colonies were dislodged from the reef framework using hammer and chisel under their dead bases and transported in buckets filled with seawater by boat to the Smithsonian Tropical Research Institute Bocas del Toro Research Station (STRI- Bocas). Collection and research were approved by the Ministerio de Ambiente de Panamá (permit no. SE/A-110-16). Colonies were placed in the outdoor flow-through experimental aquarium, which supplies seawater from Bahia del Almirante with natural temperature (29.2 ± 0.1 °C), light, and photoperiod (12:12 h). Colonies were allowed to recover and were kept in these common garden conditions for at least 10 days before being used for experiments. Internal transcribed spacer 2 (ITS2) genotyping of coral colonies taken from the same sampling site around the same time revealed *P. astreoides* predominantly hosted Symbiodiniaceae type A4 while *O. franksi* predominantly hosted Symbiodiniaceae type A3 (Linsmayer et al., 2024).

### Exposure to elevated CO_2_

The experimental exposures took place between October 26 and November 5, 2016. To generate ocean acidification-like seawater, CO_2_ was bubbled into four 200 L sumps receiving flowing seawater using a pH-stat system (IKS Aquastar, Karlsbad, Germany) that continuously measured pH and injected CO_2_ to achieve the target level (Hamilton et al., 2014; Kwan and Tresguerres, 2022; Kwan et al., 2017) (**Fig. S1**). Each sump supplied elevated CO_2_-seawater to one 45 L tank (for a total of four OA tanks), and four additional identical tanks received unmodified seawater from the STRI-Bocas aquarium system (Control tanks). The eight tanks were randomly distributed on an aquarium water table, and seawater flow was adjusted to 1 L min^-1^_._ Colonies were randomly assigned to one of the eight tanks such as each tank housed three *P. astreoides* and three *O. franksi* colonies. A submergible power head (Domica AM-005, 90 GPH, 5 watts) was used to circulate seawater in each tank.

Upon placement of the coral colonies in the tanks, the pH in the elevated CO_2_ seawater was progressively lowered to ∼7.90 (Day 1 of the exposures), then to ∼7.80 (Days 2 and 3), and finally to the target pH of ∼7.70 (Days 4-10). Throughout the exposures, seawater pH was monitored using the IKS system and additionally “spot-checked” using a handheld sensor (HACH HQ40d with pH probe PHC101) at least twice per day. Salinity (32.1 ppt) and dissolved O_2_ were measured periodically using a handheld multimeter (YSI Pro2030, Xylem Inc., Washington D.C., USA). Photosynthetically active radiation (PAR) was continuously monitored using an Odyssey PAR logger (Dataflow Systems Ltd., Christchurch, New Zealand) placed inside one of the experimental tanks. The PAR sensor was moved among tanks daily to capture the average light environment experienced by the coral colonies throughout the experiment. Because dissolved O, measurements were collected as part of routine water-quality monitoring rather than as a pre-planned light-response experiment, measurements were not conducted at predefined irradiances. Instead, PAR at the time of each O, measurement was obtained from the continuous light record and used to examine the relationship between incident irradiance and dissolved O, concentrations in control and elevated CO_2_ tanks. Tanks were cleaned daily to minimize algal growth on walls and piping of the sumps and coral tanks.

Discrete seawater samples were collected from the coral holding aquaria for pH and Total Alkalinity (TA) measurements in the Dickson laboratory (SIO, UCSD). These pH and TA values, together with temperature and salinity, were used to calculate dissolved inorganic carbon and pCO_2_ using CO2SYS (Lewis and Wallace, 1998) (**STable 1-4**). After 10 days of exposure, each coral colony was removed from the tank, photographed for colorimetric determination of pigment content, flash-frozen, and stored in liquid N_2_ and then at -80°C for antioxidant defenses and oxidative damage assays.

### O_2_ microsensor measurements

O, concentrations within the coral diffusive boundary layer (DBL) were measured as previously described (Linsmayer et al., 2024). Measurements were performed in a modified intermodal container serving as a wet laboratory, using a 45 L tank placed inside a temperature-controlled water bath maintained at 29 °C. A combination of LED and incandescent lamps suspended above the tank provided an irradiance of approximately 650 μmol photons m⁻² s⁻¹ to approximate the upper range of irradiances recorded in the outdoor experimental tanks. Irradiance was measured at the top of each colony using an Odyssey PAR logger.

O, concentrations were measured using a Clark-type microsensor with a 90% response time of less than 0.5 s and a tip diameter of 25 μm (Ox25-fast, Unisense A/S, Aarhus, Denmark). Using a manual micromanipulator and a binocular dissection microscope, the microsensor was carefully lowered onto the coenosarc tissue until a spike in O, concentration indicated contact with the coral surface. The microsensor was then slowly withdrawn until a stable O, recording was obtained, maintaining an angle of approximately 40° from vertical. Corals were allowed to adjust to the measurement conditions for 10 min before O, concentrations were recorded for an additional 10 min using Unisense SensorTrace Logger software at a sampling frequency of 1 Hz. Results represent the average O, concentration during each 10-min recording period. When seawater exchanges were required, the replacement seawater was added slowly without disturbing the microsensor position, and the exchange was completed within 2 min.

Two sets of coral colonies were used to examine acute and chronic responses to elevated CO,. For the acute experiments, a separate set of *Orbicella franksi* and *Porites astreoides* colonies was collected from the field and maintained under common-garden conditions for at least 24 h before measurements. On the first experimental day, O, concentrations were measured in control seawater before and after replacement with fresh control seawater to determine whether the exchange procedure itself affected DBL O, concentrations. On the following day, the same colonies were initially measured in control seawater and subsequently exposed to elevated-CO, seawater (∼pH 7.75). O, concentrations were recorded for 10 min under each condition using the procedure described above.

For the 10 day-exposure experiments, a subset of colonies maintained in the outdoor control and elevated-CO, treatments was transported individually to the wet laboratory on Days 8 and 9. Colonies were transported in seawater from their respective treatments and measured under the corresponding control or elevated-CO, conditions, without switching treatments. Measurements alternated between species and treatments to minimize potential effects of measurement order (N = 6 colonies per treatment).

### Antioxidant defenses and oxidative damage assays

Frozen samples were transported in a dry shipper to SIO-UCSD and stored at -80°C. Frozen coral tissue was removed from the skeleton by blasting buffer [100 mM Tris supplemented with 0.5 mM phenylmethylsulfonyl fluoride and 0.2 mM benzamidine, pH 7.5] using an airbrush (Paasche, MIL#3 Millennium Airbrush). The resulting homogenates were sonicated (four pulses of 10 sec on ice, with a 50 sec rest in between pulses), vortexed, and centrifuged (11,000 xg, 20 min, 4 °C). The resulting coral tissue-enriched supernatants (“coral sample”) were aliquoted, flash-frozen in liquid N_2_, and stored at −80°C.

Total soluble protein content was measured on all aliquots by the method of Bradford(Bradford, 1976), using bovine serum albumin as the standard. Superoxide dismutase (SOD, EC 1.15.1.1) activity was estimated based on the inhibition of nitroblue tetrazolium reduction rate (Beauchamp and Fridovich, 1971). The assay mix contained 1, 2 or 3 μL coral sample in buffer (0.1 mM EDTA, 13 mM DL-methionine, 75 μM nitroblue tetrazolium and 20 μM riboflavin, 50 mM phosphate buffer, pH 7.9, final volume 200 μL). Samples were exposed for 15 min to intense cool-white light and then kept in the dark until measuring their absorbance at 560 nm. One SOD enzymatic unit (U) was defined as the amount of enzyme required to inhibit the reaction rate by 50%. Results were expressed in U mg protein^-1^. Catalase activity (CAT, EC1.11.1.6) was measured by monitoring the decay of H_2_O_2_ at 240 nm during 30 s using an extinction coefficient of 40 M^−1^ cm^−1^ (Aebi, 1984). The assay mix contained 20 μL coral sample in buffer (50 mM potassium phosphate buffer and 10 mM H_2_O_2_, pH 7.4, final volume 1 mL). One CAT U was defined as the amount of enzyme required to transform 1 mmol H_2_O_2_ min^-1^. Results were expressed in U mg protein^-1^. Glutathione-S-transferase (GST, EC 1.11.1.9) activity was measured based on the conjugation rate of reduced glutathione (GSH) with 1-chloro-2,4-dinitrobenzene (CDNB) measured as the change in absorbance at 340 nm over 90 s (Habig et al., 1974). The assay mix contained 10 μL coral sample in buffer (1 mM GSH, 1 mM CDNB in 100 mM phosphate buffer, pH 7.5, final volume 200 μL). One GST Unit was defined as the amount of enzyme needed to catalyze the formation of 1 mmol of GS-DNB min^-1^. GSH levels were based on the oxidation of GSH by 5,5-dithiobis-(2-nitrobenzoic) acid (DTNB) and the resulting absorbance at 412 nm after 30 min incubation (Anderson, 1985). This procedure involved the deproteinization of the samples using 10 % sulfosalicylic acid. The assay mix contained 100 μL coral sample (acidified with 0.3% sulfosalicylic acid) in buffer (6 mM DTNB and 6.3 mM EDTA in 0.143 M sodium sulfate buffer, pH 7.5, final volume 1 mL). GSH levels were determined based on a standard curve generated with known GSH amounts. Results were expressed as nmol_GSH_ equivalents mg protein^-1^. Lipid peroxidation was determined measuring thiobarbituric acid reactive substances (TBARS) (Buege and Aust, 1978). The assay mix contained 175 μL coral sample in 1.4 mL thiobarbituric acid (26 mM) solution, 1.575 mL, final volume), incubated at 95°C for 45 min, cooled on ice, centrifuged (8,000 xg, 10 min), and supernatant absorbance was determined at 535 nm. TBARS concentration was estimated as malondialdehyde (MDA) equivalents, using the extinction coefficient of MDA–thiobarbituric acid complex (156 mM^−1^ cm^−1^). Results were expressed as μmol_TBARS_ mg protein^-1^. Protein carbonyl content was measured on 200 μL coral sample and a commercial kit following manufacturer’s instructions (#10005020 Cayman Chemical Company, Ann Arbor, MI). Results were expressed as nmol_carbonyl_ mg protein^-1^_._ Spectrophotometric readings were taken at 28°C (SOD, CAT, GST) or room temperature (approx. 23°C) (GSH, TBARS, carbonyl content) on a single-cell spectrophotometer (UV-1280, Shimadzu Scientific Instruments, Maryland, USA) or a plate reader SpectraMax M2 (Molecular Devices, San Jose, CA).

### Colorimetric determination of pigment content

Coral colonies were photographed using a digital camera right before starring the exposures (Day 0) and immediately prior to flash-freezing (Day 10) (**SFig. 2)**. Images were captured from a top-down perspective, maintaining consistent geometric orientation between timepoints to minimize variation due to viewing angle. During photography, each coral was positioned adjacent to the Coral Health Chart side E (Siebeck et al., 2006) to provide a standardized color reference. The square E1 of the chart was used as the white reference in all photographs.

**Figure 1.**
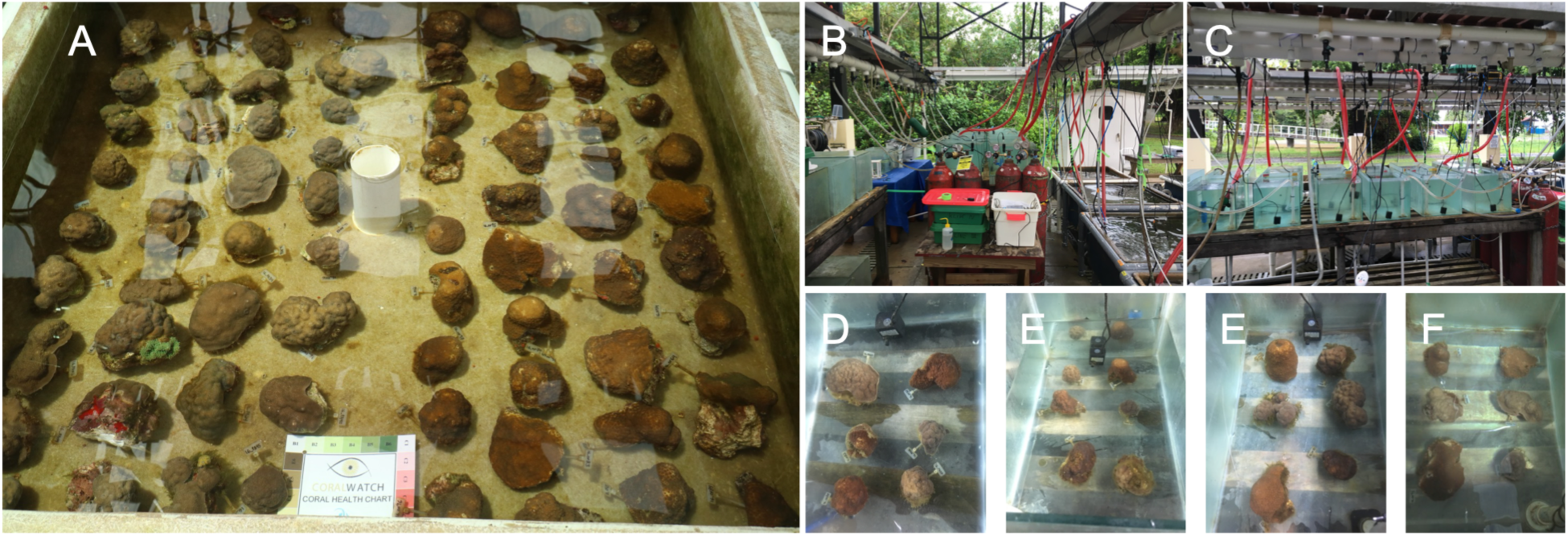
Experimental aquarium at STRI Bocas del Toro Research station. **(A)** Common garden acclimation of *Porites astreoides* and *Orbicella franksi* collected from Punta Caracol. **(B, C)** Experimental setup to manipulate CO_2_/pH levels including sumps and specimen-holding tanks. **(D-F)** Top view of specimen-holding tanks, each with three *P. astreoides* and three *O. franksi* colonies.

**Figure 2.**
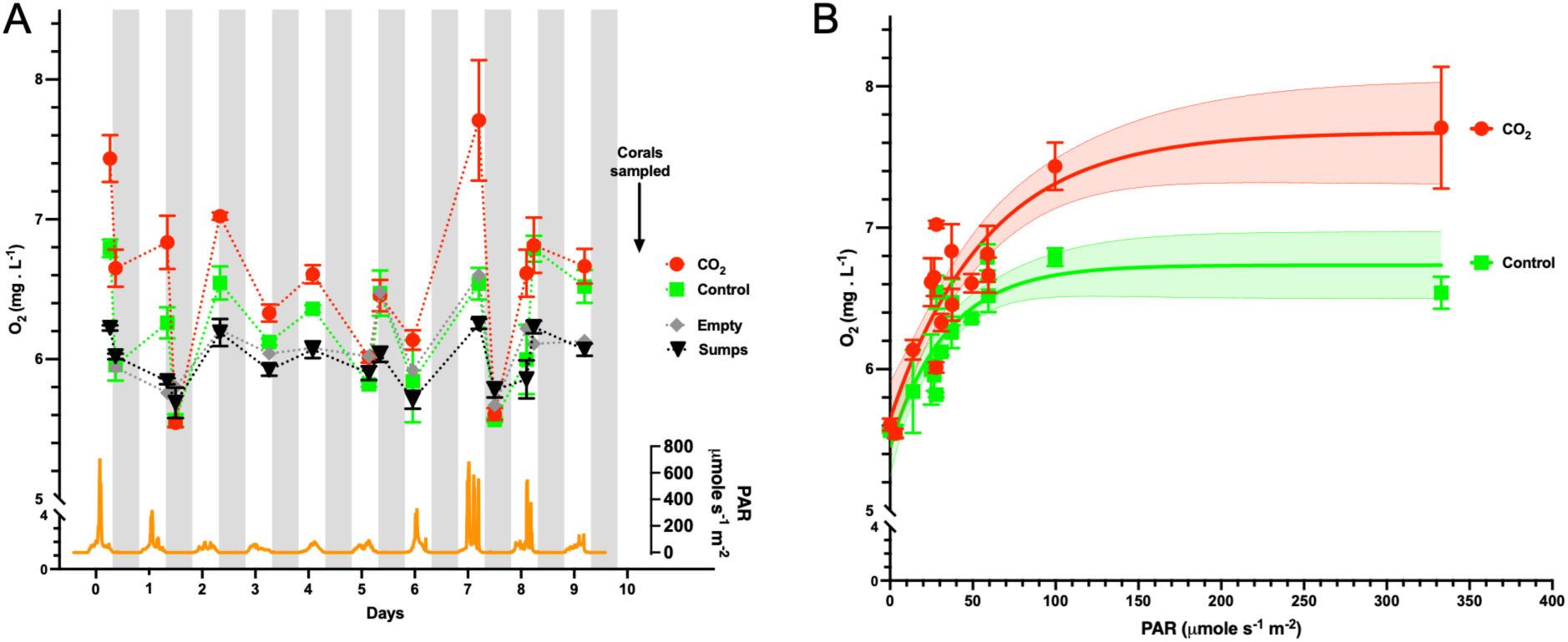
Elevated CO₂ was associated with increased dissolved O₂ concentrations in outdoor flow-through aquaria, with the largest differences occurring at higher irradiance. **(A)** Dissolved O₂ concentrations measured during routine monitoring of aquaria containing corals exposed to control (∼400 µatm) or elevated (∼1,100 µatm) CO₂, as well as empty tanks and the elevated CO_2_ seawater supply sumps. The alternating shading in the x-axis indicates successive days of the experiment. *Bottom:* Photosynthetically active radiation (PAR) recorded continuously with an Odyssey PAR sensor placed within the experimental aquaria. **(B)** Dissolved O₂ concentrations plotted against PAR measured at the corresponding time. Solid lines indicate nonlinear least-squares fits using an exponential plateau model, and dashed lines indicate 95% confidence bands. The fitted relationships differed significantly between control and elevated CO_2_ tanks (extra sum-of-squares F test, F₃,₁₁₄ = 13.81, P < 0.0001). Dissolved O₂ measurements were collected opportunistically during routine aquarium monitoring rather than at predefined irradiances and therefore do not constitute a controlled photosynthesis-irradiance experiment.

To quantify variation in coral pigmentation throughout the experiment, images were analyzed using ImageJ software (National Institutes of Health, USA). Briefly, the mean pixel intensity of the coral tissue was measured using the polygon selection tool, while the mean intensity of the white reference (E1) was measured using the rectangle selection tool (**SFig. 3**). Coral absorptance in the photosynthetically active radiation (PAR) range was calculated assuming 100% reflectance of the white reference using the following equation:

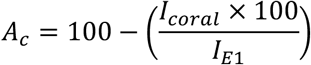

where *I_coral_*is the mean pixel intensity of the coral tissue and *I_E_*_1_is the mean pixel intensity of the white reference square. Coral absorptance (*A_c_*) was calculated for each coral at the beginning and at the end of the experiment (supplementary material photos).

**Figure 3.**
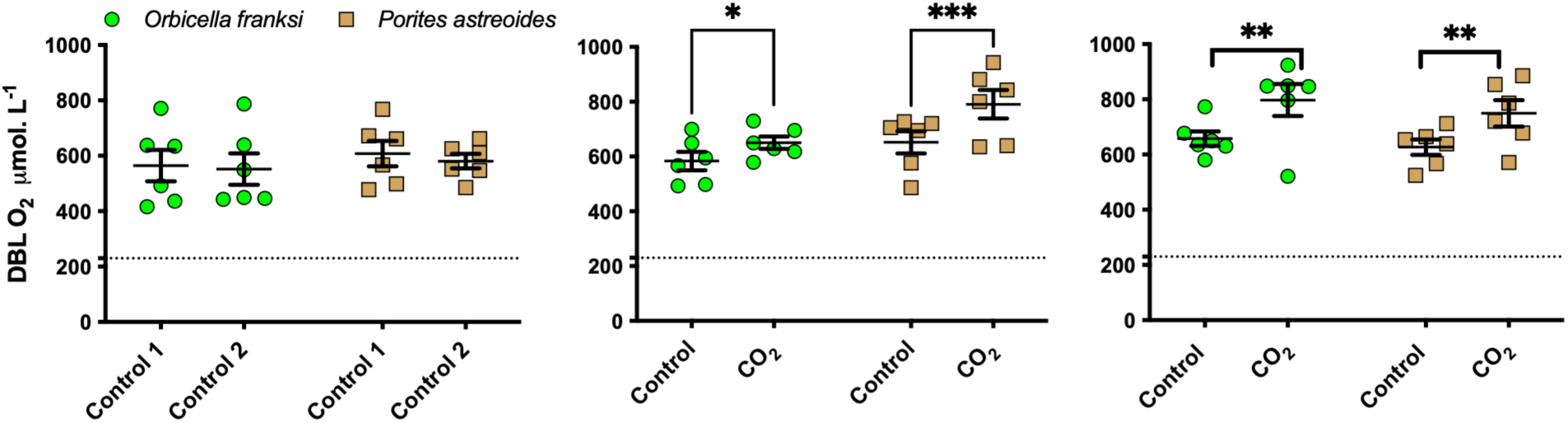
Acute and 10-day exposure to elevated CO₂ increased O₂ concentrations within the coral diffusive boundary layer. Oxygen concentrations were measured using needle microelectrodes positioned within the diffusive boundary layer (DBL) of *Orbicella franksi* and *Porites astreoides* colonies under 650 µmol photons m⁻² s⁻¹ illumination. **(A)** Replacing control seawater with fresh control seawater did not significantly change DBL O_2_ concentrations in either species. **(B)** On the following day, acute exposure of the same colonies to elevated CO_2_ significantly increased DBL O₂ concentrations in both species, with a larger response in *P. astreoides*. **(C)** Colonies from the outdoor exposure experiment were measured in seawater corresponding to their respective treatments (control or elevated CO_2_). DBL O₂ concentrations were significantly higher in elevated-CO₂-exposed colonies in both species, with similar increases in *O. franksi* and *P. astreoides*. Dotted horizontal lines indicate dissolved O_2_ concentrations in the surrounding bulk seawater. Asterisks indicate significant differences between conditions or treatments within each species (*p<0.0255, **p<0.003, ***p<0.0002; Two-way ANOVA followed by Šídák’s multiple comparisons test). Data are mean ± SEM (N = 6 colonies per treatment).

### Statistical analyses

Data is presented as mean ± SEM and analyzed using two-way analysis of variance (2-way ANOVA) followed by Šídák’s multiple comparisons tests when appropriate as described in the figure captions. The relationships between irradiance and dissolved O, concentration in control and elevated-CO, tanks were fitted by nonlinear least-squares regression using an exponential plateau model and compared using an extra sum-of-squares F test. All statistical analyses were done in Prism 11.0.2 (100) for macOS (GraphPad Software, La Jolla, CA).

## Results

### Elevated CO_2_ increased dissolved O_2_ concentrations in experimental aquaria in an irradiance-dependent manner

Routine monitoring of the experimental aquaria revealed consistently higher dissolved O_2_ concentrations in coral-holding tanks receiving elevated CO, seawater than in coral-holding tanks receiving control seawater (**Fig. 2A**). Although dissolved O, was measured primarily to monitor water quality rather than as a planned response variable, the difference between treatments became particularly apparent when measurements coincided with periods of high solar irradiance. Continuous PAR measurements confirmed that the outdoor tanks experienced large fluctuations in natural light throughout the experiment, with irradiance frequently exceeding 500–700 µmol photons m⁻² s⁻¹ during peak daylight periods (**Fig. 2B**).

Plotting dissolved O, concentrations against the PAR irradiance measured at the time of each measurement revealed that O, increased with irradiance in both treatments, but the fitted relationships differed significantly between control and OA tanks (**Fig. 2C**). The fitted curves indicated progressively higher O, concentrations under elevated CO, as irradiance increased.

Because O, measurements were collected opportunistically, most occurred at irradiances below ∼100 µmol photons m⁻² s⁻¹, with only one measurement occurring at ∼330 µmol photons m⁻² s⁻¹. Thus, the relationship should not be interpreted as a controlled photosynthesis-irradiance response curve. Nevertheless, the consistent treatment difference and its dependence on irradiance suggested that elevated CO, enhanced net photosynthetic O, production. These observations motivated experiments specifically designed to test this hypothesis by measuring O, directly within the coral diffusive boundary layer under controlled irradiance.

### Acute and chronic exposure to elevated CO, enhanced photosynthetic O, production

To determine whether elevated CO, directly altered photosynthetic O, production, microelectrodes were positioned within the diffusive boundary layer (DBL) overlying colonies of *P. astreoides* and *O. franksi* (**Fig. 3**). Based on the irradiances recorded in the outdoor aquaria, microelectrode experiments were conducted at ∼650 µmol photons m⁻² s⁻¹ to approximate the higher light levels experienced by the corals during the exposure experiment. Oxygen concentrations within the DBL substantially exceeded those in the surrounding bulk seawater, reflecting the strong metabolic modification of the coral surface microenvironment during photosynthesis. To determine whether seawater exchange itself affected DBL O, concentrations, control seawater was replaced with fresh control seawater without disturbing the microsensor position. This procedure did not significantly alter DBL O, concentrations in either species (**Fig. 3A**). Acute exposure of the same colonies on the following day to elevated CO, significantly increased DBL oxygen concentrations in both species (**Fig. 3B)**. This response was particularly pronounced in *P. astreoides*, where oxygen concentrations increased from 651.7 ± 40.0 to 790.5 ± 52.0 µmol L⁻¹, whereas *O. franksi* exhibited a smaller increase from 583.5 ± 33.6 to 650.3 ± 22.3 µmol L⁻¹. In colonies from the outdoor exposure experiment, measured in seawater corresponding to their respective treatments, both species exhibited significantly higher O, concentrations within the DBL than under control conditions. In *P. astreoides*, the difference between control and elevated CO_2_ water (657.0 ± 26.4 vs. 797.5 ± 57.6 µmol L⁻¹) was nearly identical to that observed during the acute exposure. In contrast, *O. franksi* exhibited a larger difference between treatments (626.7 ± 28.0 vs. 749.2 ± 47.7 µmol L⁻¹), consistent with enhanced photosynthetic oxygen production following chronic exposure to elevated CO,.

### 10-day exposure to elevated CO_2_ did not alter coral pigmentation

To determine whether enhanced photosynthetic O, production was associated with changes in algal abundance or photopigment content, coral coloration was quantified from standardized photographs using image analysis (**Fig. 4**). Neither *P. astreoides* nor *O. franksi* exhibited significant changes in tissue absorptance after 10 days of exposure to either control or elevated CO, conditions. Similar absorptance values at the beginning and end of the experiment suggest that elevated CO_2_ did not induce detectable changes in overall pigmentation during the experimental period. Although absorptance measurements cannot distinguish between changes in Symbiodiniaceae density and changes in pigment concentration per cell, the absence of detectable color changes indicates that the enhanced O, production observed under elevated CO, was unlikely to result from large shifts in symbiont abundance or chlorophyll content over this relatively short exposure.

**Figure 4.**
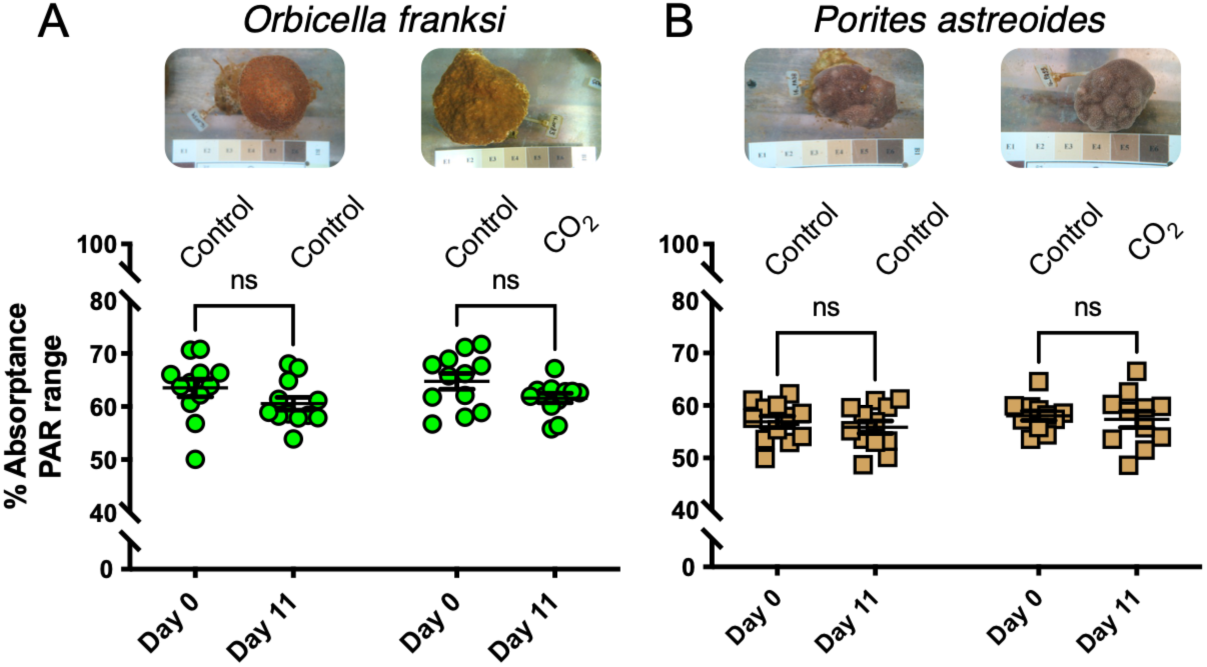
Elevated CO_2_ did not alter coral tissue pigmentation after 10 days of exposure. Coral tissue absorptance in the photosynthetically active radiation (PAR) range was quantified from standardized digital photographs as described in the Methods. Neither *Orbicella franksi* **(A)** nor *Porites astreoides* **(B)** exhibited significant changes in tissue absorptance between Day 0 and Day 10 under control or elevated CO₂ conditions. Photographs show representative control and elevated CO_2_ colonies after the 10-day exposure. Tissue coloration is commonly used as a proxy for Symbiodiniaceae abundance and photosynthetic pigment content; these results suggest that the enhanced O_2_ production following chronic exposure to elevated CO_2_ shown in Figure 3C was not accompanied by large changes in either parameter. Boxes represent the interquartile range, center lines the median, whiskers the minimum and maximum values, and symbols individual colonies (N = 6).

### Antioxidant defenses remained stable despite enhanced photosynthetic oxygen production

To determine whether increased photosynthetic O, production under elevated CO, resulted in oxidative stress, we quantified multiple components of the antioxidant defense system together with markers of oxidative damage on corals exposed to control or elevated CO_2_ for 10 days (**Fig. 5**). Exposure to elevated CO, did not significantly affect the activities of SOD, catalase, or glutathione-S-transferase in either coral species. Likewise, reduced glutathione concentrations remained unchanged between treatments. Lipid peroxidation, assessed as TBARS, was also unaffected by elevated CO, in both species. Protein oxidation exhibited a species-specific response. Carbonyl content remained unchanged in *O. franksi* but was significantly lower in *P. astreoides* following exposure to elevated CO,. Independent of treatment, the total levels of several antioxidant parameters differed between species, indicating inherent physiological differences in redox metabolism between *P. astreoides* and *O. franksi*.

**Figure 5.**
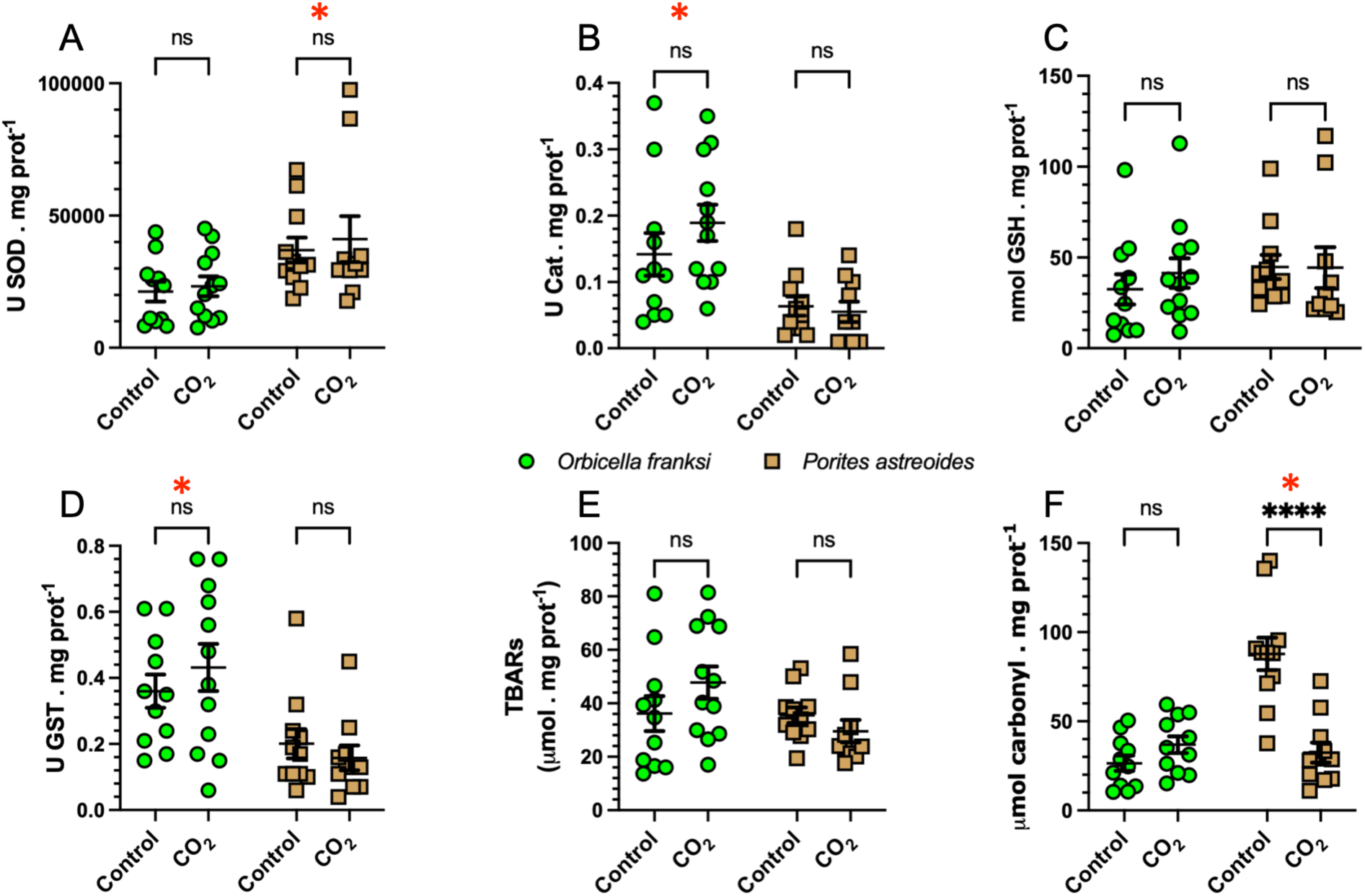
Elevated CO_2_ had limited effects on antioxidant defenses and oxidative damage but decreased protein oxidation in *Porites astreoides*. Activities of superoxide dismutase (SOD), catalase (CAT), glutathione-S-transferase (GST), reduced glutathione (GSH), lipid peroxidation (TBARS), and protein carbonyl content were measured after 10 days of exposure to control or elevated CO₂ conditions. Ocean acidification did not significantly alter antioxidant enzyme activities, glutathione levels, or lipid peroxidation in either species. Likewise, protein carbonyl content remained unchanged in *Orbicella franksi* but was significantly reduced in *Porites astreoides*. A red asterisk indicates significant differences between species irrespective of treatment, whereas the four black asterisks indicate significant effects of elevated CO₂ within a species (p<0.0001; Two-way ANOVA followed by Šídák’s multiple comparisons test). Data are mean ± SEM (N = 10–12 colonies per treatment).

## Discussion

Acute and 10-day exposure to elevated CO, increased photosynthetic oxygen production in both *P. astreoides* and *O. franksi*, resulting in higher O, concentrations both within the coral DBL and in bulk mesocosm seawater during daylight hours. Despite this increase in O_2_ production, neither species exhibited evidence of increased oxidative stress or activation of antioxidant defenses. Instead, *P. astreoides* experienced decreased protein oxidation under elevated CO,, suggesting that improved inorganic carbon availability may have enhanced photosynthetic efficiency while decreasing oxidative damage. Together, these results indicate that elevated CO, can simultaneously stimulate photosynthetic oxygen production and maintain, or even improve, cellular redox homeostasis, while raising the possibility that future ocean acidification may influence reef oxygen dynamics.

### Elevated CO_2_ increased O_2_ concentrations in the aquaria, particularly at higher irradiance

The consistently higher dissolved O_2_ concentrations in the outdoor flow-through aquaria exposed to elevated CO, were unexpected. Although O_2_ measurements were initially collected as part of routine monitoring of the experimental system rather than as a planned response variable, O_2_ concentrations were repeatedly higher in the elevated CO, treatment. Moreover, the relationship between irradiance and O, concentration differed significantly between control and elevated CO, tanks, with the fitted curves showing progressively higher O, concentrations under elevated CO, as irradiance increased.

Nevertheless, these observations should be interpreted cautiously. The experiment was not designed as a controlled light-response experiment, and dissolved O, was measured opportunistically at the irradiance occurring when routine aquarium monitoring was performed. Consequently, most O, measurements occurred under relatively low to moderate irradiance even though PAR measurements showed that the corals regularly experienced substantially higher irradiances. In addition, although tanks were cleaned daily and coral colonies were randomly distributed among treatments, we cannot completely exclude contributions from photosynthetic organisms growing on tank surfaces or small differences in the combined biomass or photosynthetic capacity of the colonies within individual tanks. Importantly, however, these potential sources of O, were exposed to the same experimental CO, manipulation and therefore do not negate the observation that the elevated CO, aquaria developed higher daytime O, concentrations.

The irradiance range represented by the O, measurements also suggests that the maximum treatment differences were probably not captured. Most measurements were made below ∼100 µmol photons m⁻² s⁻¹ and only one occurred near 330 µmol photons m⁻² s⁻¹, whereas PAR periodically exceeded 700 µmol photons m⁻² s⁻¹ around noon. We therefore cannot determine how high dissolved O, concentrations became during periods of maximum irradiance. This observation was one reason we subsequently performed the coral microsensor experiments at ∼650 µmol photons m⁻² s⁻¹, approximating the higher irradiances experienced by the colonies in the outdoor tanks.

This pattern is consistent with elevated CO, stimulating photosynthetic O_2_ production by the coral colonies, and with the additional inorganic carbon becoming increasingly important as photosynthetic activity increased with light availability. This finding is noteworthy because elevated CO, is commonly associated with low O_2_ coastal marine ecosystems (Chan et al., 2019; Gobler et al., 2014; Klein et al., 2020; Pörtner et al., 2005; Steckbauer et al., 2020). However, this relationship largely reflects biological respiration, which consumes O, while producing CO,. In contrast, OA is driven by the absorption of atmospheric CO, into surface seawater and therefore does not inherently decrease O, availability. Instead, if photosynthetic organisms are carbon limited under present-day conditions, increased dissolved inorganic carbon may stimulate photosynthesis and increase O, production during daylight hours.

Natural shallow-water reefs can experience irradiances substantially greater than those experienced during our O, measurements, raising the possibility that CO,-dependent stimulation of photosynthesis could become even more pronounced under high light conditions. However, extrapolation beyond the irradiances tested here is difficult because photosynthesis eventually becomes light-saturated and very high irradiance can induce photoinhibition. Increased inorganic carbon availability could potentially modify these responses by providing additional substrate for carbon fixation as photon input increases, but this possibility cannot be resolved from the present experiment. Thus, while our aquaria measurements lack the temporal resolution required to quantify whole-system O, budgets, they raise the possibility that ocean acidification could alter daytime O, dynamics at spatial scales extending beyond individual coral colonies.

### Enhanced oxygen production extended from the coral boundary layer to the surrounding seawater

The O, microsensor measurements confirmed that the higher dissolved O, concentrations observed in the outdoor aquaria originated, at least in part, from increased photosynthetic O, production by the corals themselves. Acute exposure demonstrated that both species rapidly responded to elevated CO, with increased photosynthetic O, production, indicating that CO_2_ availability can immediately influence photosynthetic performance. Chronic exposure maintained this response in *P. astreoides* and further enhanced it in *O. franksi*, suggesting that the two species differ in the timescale over which they physiologically adjust to elevated CO,. Coral pigmentation did not change following the 10-day exposure to elevated CO, (**Fig. 3**), suggesting that the enhanced photosynthetic O, production was not driven by major changes in symbiont abundance or chlorophyll content (Anthony et al., 2008; Chow et al., 2016; Cooper and Fabricius, 2012; Winters et al., 2009). However, these parameters could not be measured directly, so contributions from changes in symbiont abundance or chlorophyll content to the enhanced photosynthetic O, production under elevated CO, cannot be excluded. Such changes would not alter our central conclusions, but future studies could determine the relative contributions of increased dissolved inorganic carbon availability and potential longer-term photoacclimation of the algal symbionts. Importantly, the rapid increase in O, production during acute exposure is more consistent with enhanced DIC availability than with changes in symbiont abundance or pigment content.

Together, the aquaria and microsensor measurements imply that the effects of elevated CO_2_ on coral O, production extended from the coral tissue interface to the surrounding water column. If elevated CO, similarly stimulates photosynthetic O, production in reef-building corals and other benthic primary producers in natural coral reefs, these could experience higher daytime dissolved O, concentrations despite lower seawater pH. However, the extent to which this phenomenon influences O, dynamics at the scale of entire reef communities remains uncertain and will depend on community composition, irradiance, water motion, and the balance between photosynthesis and respiration. Whether similar responses occur in other reef primary producers and how they influence diel O, dynamics in natural reef communities remain important questions for future investigation.

### Enhanced photosynthetic oxygen production did not result in increased oxidative stress

An increase in photosynthetic O, production might be expected to increase the generation of ROS leading to oxidative damage (Downs et al., 2002; Lesser, 2006; Morris et al., 2022; Szabó et al., 2020). However, our biochemical analyses did not support that possibility. Activities of the antioxidant enzymes SOD, catalase, and GST remained unchanged following OA exposure, as did reduced glutathione concentrations and lipid peroxidation levels. These results indicate that the increase in O, production in the coral DBL was not accompanied by measurable disruption of cellular redox homeostasis during the 10-day exposure.

Notably, O, concentration alone does not determine oxidative stress. In photosynthetic organisms, ROS generation is controlled by the balance between light harvesting and downstream carbon fixation. When the Calvin cycle is limited by inorganic carbon availability, excess excitation energy and reducing equivalents cannot be fully utilized for carbon fixation and may instead be diverted toward O, production, increasing ROS production through processes such as the Mehler reaction and photorespiration (Krueger et al., 2014; Roberty et al., 2014). Under these conditions, increasing inorganic carbon availability can improve the coupling between light capture and carbon fixation, decreasing rather than increasing ROS production despite higher rates of photosynthetic oxygen evolution.

Our results are consistent with this interpretation. Our 10-day exposure to elevated CO, increased photosynthetic O, production while producing no evidence of enhanced oxidative damage. Rather than stimulating antioxidant defenses to compensate for increased ROS production, corals maintained antioxidant enzyme activities and glutathione levels similar to those observed under present-day CO, conditions. However, the different acute and chronic photosynthetic responses of the two species indicate that physiological responses to elevated CO, can develop over different timescales. Thus, transient changes in antioxidant defenses during the exposure, or responses developing over longer time scales, cannot be excluded from our endpoint measurements. These findings suggest that elevated CO, did not impose additional oxidative stress during the experimental period and instead may have improved the efficiency with which absorbed light energy was utilized for carbon fixation. This interpretation is further supported by the species-specific response of protein oxidation. Whereas *O. franksi* showed no detectable change in protein carbonyl content, *P. astreoides* exhibited significantly lower protein oxidation following exposure to elevated CO,. Protein carbonylation represents an irreversible oxidative modification that integrates oxidative damage over longer timescales than transient antioxidant responses. Consequently, the decrease in protein oxidation observed in *P. astreoides* suggests that elevated CO, not only failed to increase oxidative stress but may have decreased oxidative damage in this species.

### Decreased protein oxidation in P. astreoides is consistent with relief of carbon limitation

Although both coral species exhibited increased photosynthetic O, production following a 10- day exposure to elevated CO,, only *Porites astreoides* showed a significant decrease in protein oxidation. Interestingly, the two species differed in the temporal dynamics of their photosynthetic responses. *P. astreoides* exhibited a robust increase in O, production immediately following acute CO, exposure that was maintained after chronic acclimation, whereas *O. franksi* displayed a smaller acute response but a larger increase after 10 days of exposure. Despite these differences in acclimation dynamics, only *P. astreoides* exhibited decreased oxidative damage, suggesting that stimulation of photosynthesis alone is insufficient to explain the species-specific biochemical responses.

One possible explanation is that the symbiotic algae of *P. astreoides* were carbon limited under present-day seawater conditions and that elevated CO, partially relieved this limitation by improving the coupling between light capture and carbon assimilation, decreasing ROS production even as photosynthetic oxygen evolution increases, as previously hypothesized (Wooldridge, 2010; Wooldridge, 2013). Importantly, activities of SOD, catalase, glutathione-S-transferase, and glutathione concentrations remained unchanged throughout the experiment, indicating that the decrease in protein oxidation was not associated with upregulation of the antioxidant systems measured here. Instead, our results are more consistent with a decrease in ROS production resulting from improved coupling between light capture and carbon fixation under elevated CO,. However, additional experiments directly quantifying carbon fixation, photosynthetic electron transport, and ROS production will be required to test this idea.

The apparent relief of carbon limitation in *P. astreoides* may help explain the remarkable resilience of this species reported in previous studies. Over the past two decades, *P. astreoides* has repeatedly emerged as one of the most environmentally tolerant Caribbean corals, maintaining comparatively high physiological performance under elevated CO_2_, elevated temperature, bleaching events, and other environmental disturbances. Experimental studies have consistently shown that *P. astreoides* performs as well as or better than many other Caribbean reef builders under projected future climate conditions (Bove et al., 2019; Enochs et al., 2016), while broader meta analyses have highlighted this species as one of the clearest examples of a relatively climate-resilient Caribbean coral (Lima et al., 2022). At the community level, demographic studies have also documented an increasing relative abundance of *P. astreoides* on many Caribbean reefs as several historically dominant framework-building species such as *Acropora* have declined (Cramer et al., 2020), leading to its designation as a potential "winner species" under ongoing climate change (Darling et al., 2012; Green et al., 2008; Lima et al., 2022). In conclusion, our study demonstrates that elevated CO_2_ increased photosynthetic O, production in two Caribbean reef-building corals without inducing oxidative stress, and that *P. astreoides* experienced decreased protein oxidation under elevated CO,. These findings are consistent with the hypothesis that relief of carbon limitation can improve photosynthetic performance while decreasing oxidative damage, providing a potential physiological mechanism contributing to the resilience of this species under future ocean conditions. More broadly, our results suggest that ocean acidification has the potential to modify daytime O, dynamics from the coral surface to the surrounding reef environment, emphasizing that the biological consequences of elevated CO, cannot be inferred from seawater chemistry alone. Integrating measurements across multiple levels of biological organization, from microscale O, gradients to whole-organism physiology and reef-scale processes, will be essential for developing a more mechanistic understanding of coral responses to ongoing ocean change.

## Acknowledgements

This study was supported by National Science Foundation Grant OCE 1538495 to DIK and MT, and an Alfred P. Sloan Research Fellowship (BR2013-103) and UC San Diego-SIO discretionary funds to MT. LBL was supported by a STRI predoctoral fellowship hosted by Dr. Nancy Knowlton and Dr. Andrew Altieri (STRI). CTGM was supported by National Science Foundation Grant IOS 2149926 to MT. ABT was supported by the National Science Foundation Graduate Research Fellowship Program (2019271478). SES received a Travelling Fellowship from the Company of Biologists to visit and conduct experiments at the Tresguerres Laboratory (SIO-UCSD). We are grateful to the staff from the Bocas del Toro Research station (STRI) for fieldwork support, especially Plinio Góndola, Deyvis González, and Dr. Rachel Collin.

## Use of AI tools

ChatGPT (GPT-5.6 Sol; OpenAI) was used to improve human-generated texts for readability and style.

---

**Supplementary Figure 1.**
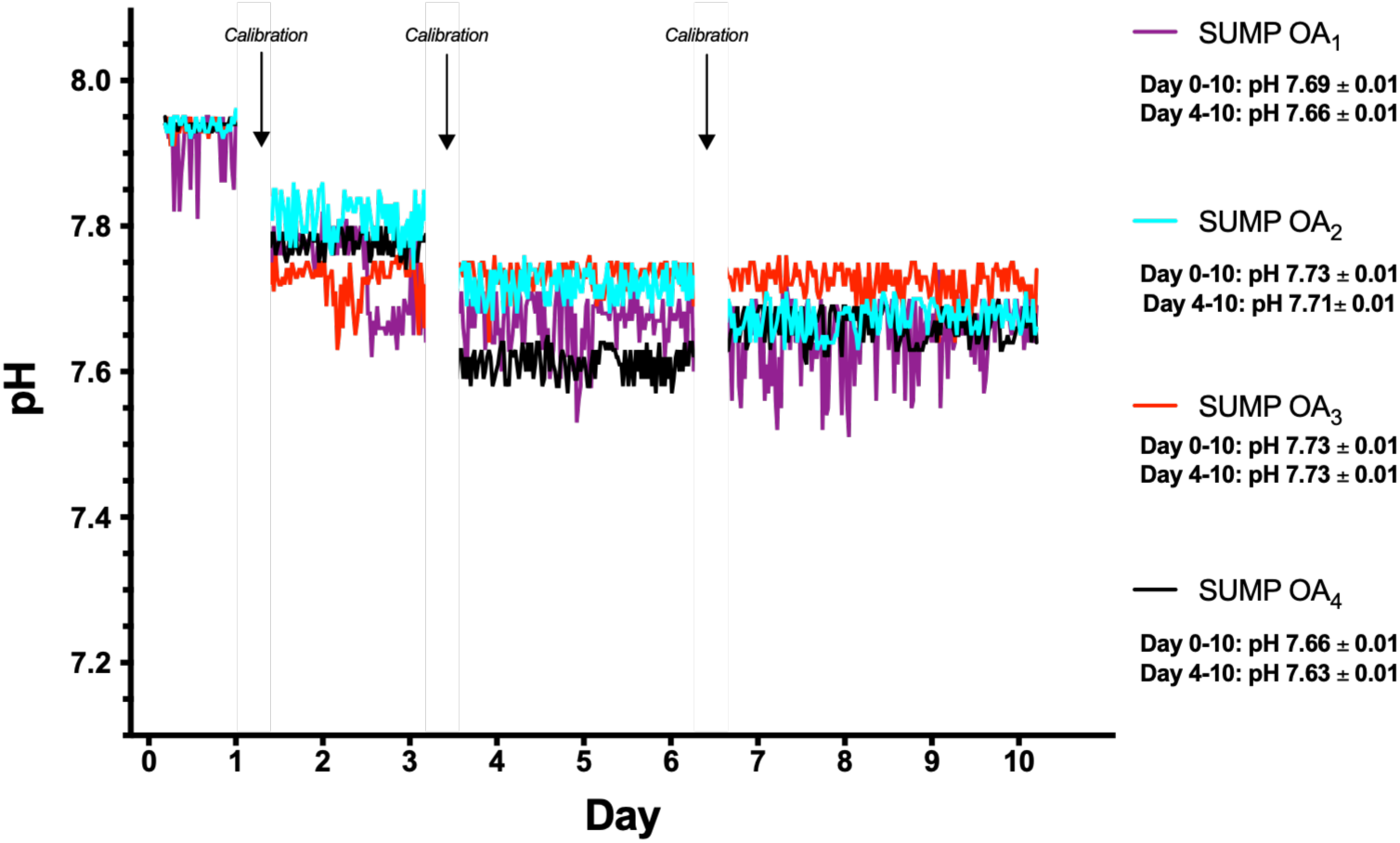
Seawater pH in the experimental OA sumps, which was monitored and recorded every 2 minutes using a pH-stat system (see Methods for more details). The gaps in the traces reflect pauses in pH recording during pH electrode calibration. Average pH in each sump (mean ± S.E.M) are shown during the entire Day 0-10 period as well as during Day 4-10 when the final target pH was achieved.

**Supplementary Figure 2.**
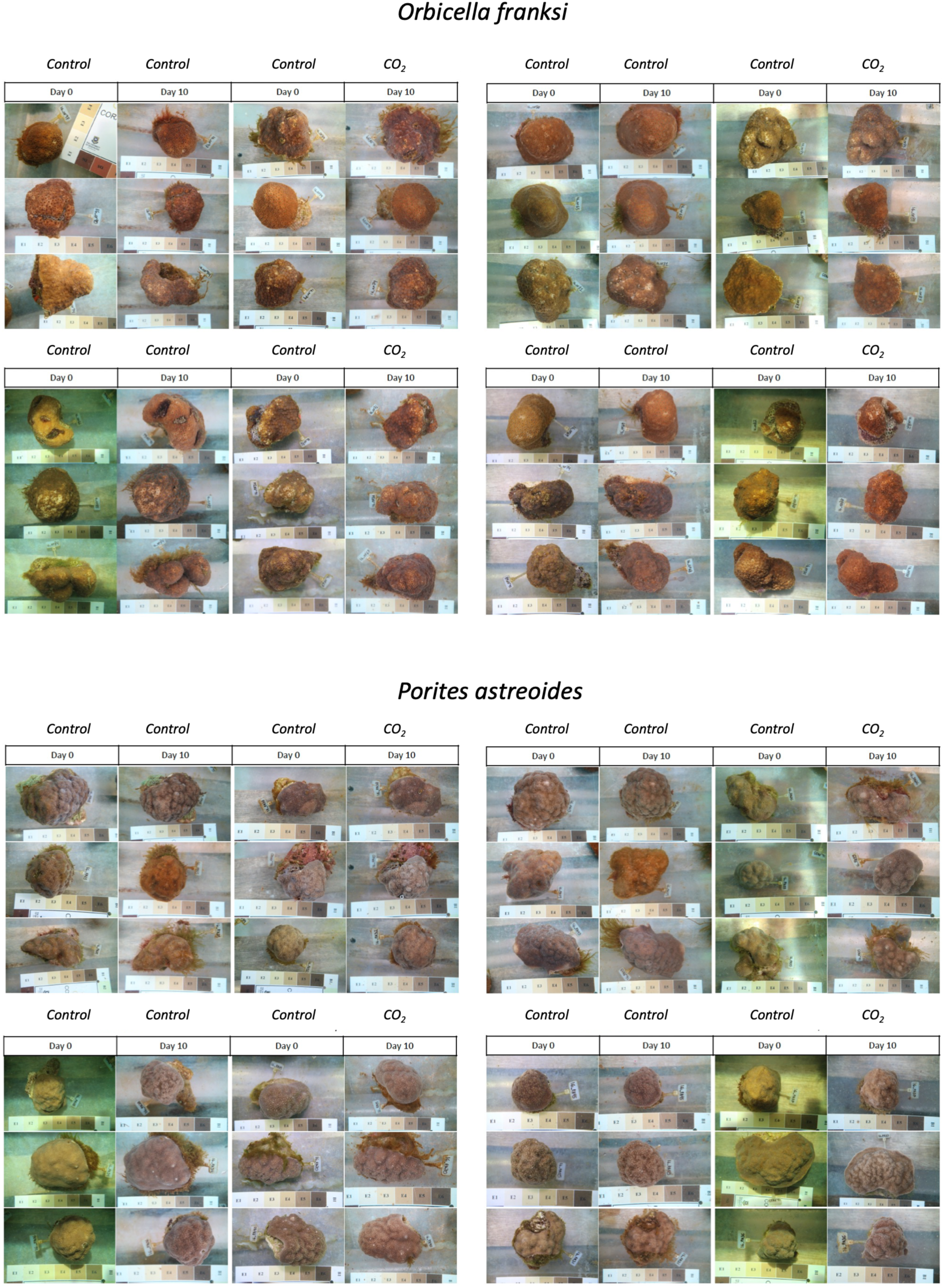
Photographs of *Orbicella franksi* and *Porites astreoides* colonies used for colorimetric determination of pigment content. See SFig. 3 for a graphical representation of the methods and Fig. 4 for the results.

**Supplementary Figure 3.**
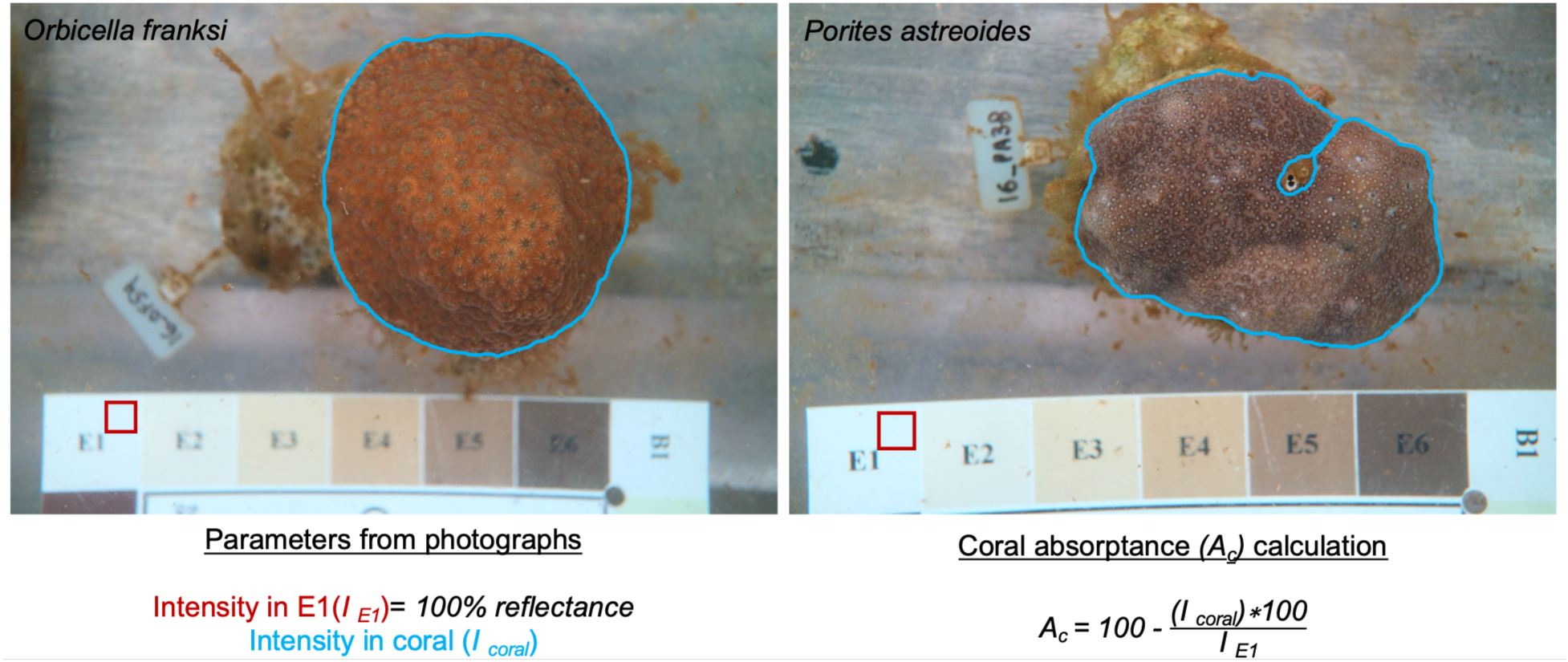
Colorimetric determination of pigment content in *Orbicella franksi* and *Porites astreoides*. Blue tracing surrounds the coral area of interest in each picture. The strip with the color squares at the bottom of each image is the Coral ColorWatch scale utilized as reference; the red square indicates 100% reflectance. Tags on the left side of each coral were used to track individual specimens.

**Supplementary Table 1.** Seawater pH measurements in discrete samples collected from the experimental tanks throughout the exposures. Tank and Treatment values are Mean ± S.E.M. Continuous pH measurements in the sumps are shown in **Fig.S2**. Other chemistry parameters are shown in **STables 2-4**.

| Control (Ctrol) |  |  |  |  | Elevated CO<sub>2</sub> |  |  |  |
| --- | --- | --- | --- | --- | --- | --- | --- | --- |
|  | Ctrol<sub>1</sub> | Ctrol<sub>2</sub> | Ctrol<sub>3</sub> | Ctrol<sub>4</sub> | OA<sub>1</sub> | OA<sub>2</sub> | OA<sub>3</sub> | OA<sub>4</sub> |
| Day 5 | 8.06 | - | - | 8.06 | 7.71 | 7.72 | 7.70 | 7.66 |
| Day 6 | - | 8.00 | 8.01 | - | 7.64 | 7.68 | 7.65 | 7.61 |
| Day 10 | 8.04 | 8.01 | 8.06 | 8.02 | 7.63 | 7.61 | 7.64 | 7.66 |
| Tank | 8.05 ± 0.01 | 8.05 ± 0.01 | 8.04 ± 0.03 | 8.04 ± 0.02 | 7.66 ± 0.03 | 7.67 ± 0.03 | 7.66 ± 0.02 | 7.64 ± 0.02 |
| Treatment | 8.03 ± 0.01 | | | | 7.66 ± 0.01 | | | |

**Supplementary Table 2.** Seawater total alkalinity (μmol.kg^-1^) measurements in discrete samples collected from the experimental tanks throughout the exposures. Tank and Treatment values are Mean ± S.E.M. Other chemistry parameters are shown in **Stables 1, 3, and 4**.

| Control (Ctrol) |  |  |  |  | Elevated CO<sub>2</sub> |  |  |  |
| --- | --- | --- | --- | --- | --- | --- | --- | --- |
|  | Ctrol<sub>1</sub> | Ctrol<sub>2</sub> | Ctrol<sub>3</sub> | Ctrol<sub>4</sub> | OA<sub>1</sub> | OA<sub>2</sub> | OA<sub>3</sub> | OA<sub>4</sub> |
| Day 5 | 2194.5 | - | - | 2192.2 | 2196.0 | 2196.0 | 2197.9 | 2196.1 |
| Day 6 | - | 2199.1 | 2194.6 | - | 2194.6 | 2199.8 | 2198.3 | 2204.7 |
| Day 10 | 2204.8 | 2198.2 | 2199.2 | 2206.0 | 2211.9 | 2203.5 | 2203.5 | 2206.8 |
| Tank | 2199.6 ± 3.7 | 2198.7 ± 0.3 | 2196.9 ± 1.6 | 2020.1 ± 4.9 | 2202.9 ± 4.1 | 2198.8 ± 2.6 | 2202.9 ± 4.2 | 2202.5 ± 2.8 |
| Treatment | 2198.6 ± 2.5 | | | | 2201.8 ± 1.8 | | | |

**Supplementary Table 3.** Calculated seawater dissolved inorganic carbon (μmol.kg^-1^) in discrete samples collected from the experimental tanks throughout the exposures. Tank and Treatment values are Mean ± S.E.M. Other chemistry parameters are shown in **Stables 1, 2, and 4**.

| Control (Ctrol) |  |  |  |  | Elevated CO<sub>2</sub> |  |  |  |
| --- | --- | --- | --- | --- | --- | --- | --- | --- |
|  | Ctrol<sub>1</sub> | Ctrol<sub>2</sub> | Ctrol<sub>3</sub> | Ctrol<sub>4</sub> | OA<sub>1</sub> | OA<sub>2</sub> | OA<sub>3</sub> | OA<sub>4</sub> |
| Day 5 | 1909.7 | - | - | 1904.2 | 2072.4 | 2066.9 | 2076.9 | 2090.2 |
| Day 6 | - | 1946.2 | 1933.5 | - | 2101.1 | 2087.6 | 2096.9 | 2116.6 |
| Day 10 | 1925.0 | 1935.4 | 1910.2 | 1936.9 | 2116.8 | 2113.7 | 2111.0 | 2098.7 |
| Tank | 1917.3 ± 5.4 | 1940.8 ± 3.8 | 1921.9 ± 8.2 | 1920.6 ± 11.6 | 2096.8 ± 11.3 | 2089.4 ± 11.7 | 2094.9 ± 8.5 | 2101.8 ± 6.7 |
| Treatment | 1925.1 ± 7.7 | | | | 2095.7 ± 5.0 | | | |

**Supplementary Table 4.** Calculated seawater CO_2_ measurements (μatm) in discrete samples collected from the experimental tanks throughout the exposures. Tank and Treatment values are Mean ± S.E.M. Other seawater chemistry parameters are shown in **Stables 1-3**.

|  | **Control (Ctrol)** |  |  |  | **Elevated CO<sub>2</sub>** |  |  |  |
| --- | --- | --- | --- | --- | --- | --- | --- | --- |
|  | **Ctrol<sub>1</sub>** | **Ctrol<sub>2</sub>** | **Ctrol<sub>3</sub>** | **Ctrol<sub>4</sub>** | **OA<sub>1</sub>** | **OA<sub>2</sub>** | **OA<sub>3</sub>** | **OA<sub>4</sub>** |
| **Day 5** | 362.9 | - | - | 356.4 | 926.6 | 908.5 | 945.0 | 1048.4 |
| **Day 6** | - | 429.4 | 409.9 | - | 1097.3 | 1004.4 | 1083.2 | 1197.9 |
| **Day 10** | 378.1 | 409.1 | 358.4 | 399.7 | 1147.6 | 1185.4 | 1095.9 | 1043.2 |
| **Tank** | 370.5 ±<br>5.4 | 419.3 ±<br>7.2 | 384.2 ±<br>18.2 | 378.1 ±<br>15.3 | 1057.2<br>± 57.9 | 1032.8<br>± 70.3 | 1041.4<br>± 41.8 | 1096.5<br>± 43.9 |
| **Treatment** | **388.0 ± 13.4** |  |  |  | **1057.0 ± 28.0** |  |  |  |

## Data Deposition

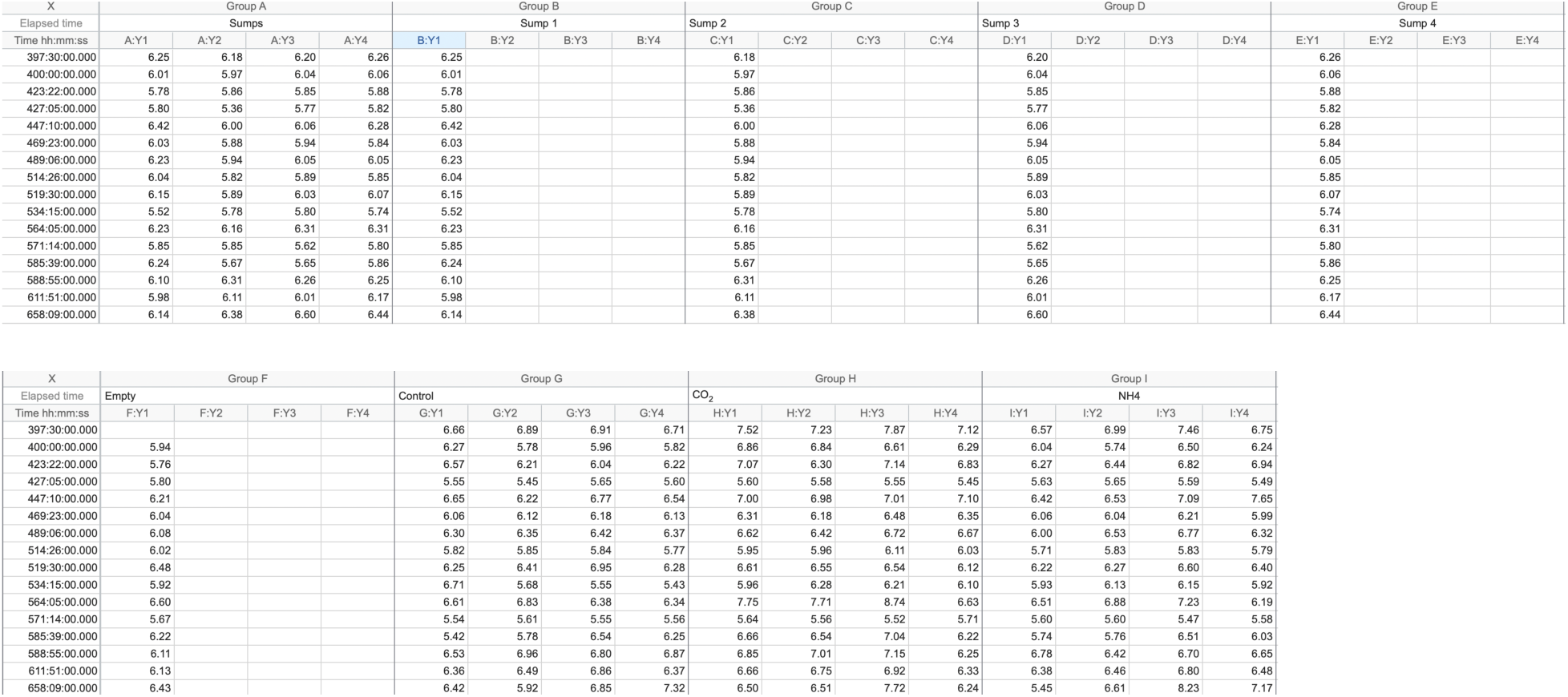

Data associated with Figure 2A.

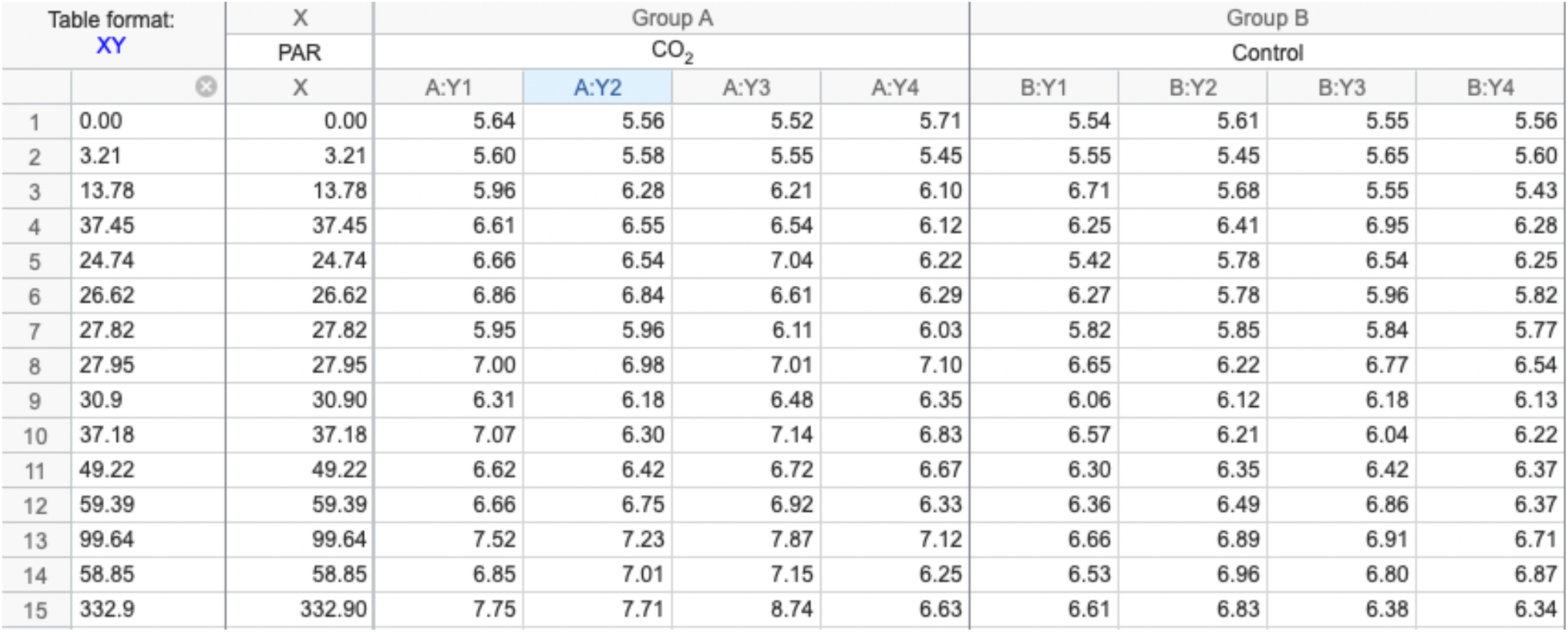

| Table format: XY |  | X | Group A |  |  |  | Group B |  |  |  |
| --- | --- | --- | --- | --- | --- | --- | --- | --- | --- | --- |
|  |  | PAR | CO<sub>2</sub> |  |  |  | Control |  |  |  |
|  | ⊗ | X | A:Y1 | A:Y2 | A:Y3 | A:Y4 | B:Y1 | B:Y2 | B:Y3 | B:Y4 |
| 1 | 0.00 | 0.00 | 5.64 | 5.56 | 5.52 | 5.71 | 5.54 | 5.61 | 5.55 | 5.56 |
| 2 | 3.21 | 3.21 | 5.60 | 5.58 | 5.55 | 5.45 | 5.55 | 5.45 | 5.65 | 5.60 |
| 3 | 13.78 | 13.78 | 5.96 | 6.28 | 6.21 | 6.10 | 6.71 | 5.68 | 5.55 | 5.43 |
| 4 | 37.45 | 37.45 | 6.61 | 6.55 | 6.54 | 6.12 | 6.25 | 6.41 | 6.95 | 6.28 |
| 5 | 24.74 | 24.74 | 6.66 | 6.54 | 7.04 | 6.22 | 5.42 | 5.78 | 6.54 | 6.25 |
| 6 | 26.62 | 26.62 | 6.86 | 6.84 | 6.61 | 6.29 | 6.27 | 5.78 | 5.96 | 5.82 |
| 7 | 27.82 | 27.82 | 5.95 | 5.96 | 6.11 | 6.03 | 5.82 | 5.85 | 5.84 | 5.77 |
| 8 | 27.95 | 27.95 | 7.00 | 6.98 | 7.01 | 7.10 | 6.65 | 6.22 | 6.77 | 6.54 |
| 9 | 30.9 | 30.90 | 6.31 | 6.18 | 6.48 | 6.35 | 6.06 | 6.12 | 6.18 | 6.13 |
| 10 | 37.18 | 37.18 | 7.07 | 6.30 | 7.14 | 6.83 | 6.57 | 6.21 | 6.04 | 6.22 |
| 11 | 49.22 | 49.22 | 6.62 | 6.42 | 6.72 | 6.67 | 6.30 | 6.35 | 6.42 | 6.37 |
| 12 | 59.39 | 59.39 | 6.66 | 6.75 | 6.92 | 6.33 | 6.36 | 6.49 | 6.86 | 6.37 |
| 13 | 99.64 | 99.64 | 7.52 | 7.23 | 7.87 | 7.12 | 6.66 | 6.89 | 6.91 | 6.71 |
| 14 | 58.85 | 58.85 | 6.85 | 7.01 | 7.15 | 6.25 | 6.53 | 6.96 | 6.80 | 6.87 |
| 15 | 332.9 | 332.90 | 7.75 | 7.71 | 8.74 | 6.63 | 6.61 | 6.83 | 6.38 | 6.34 |

Data associated with Figure 2B.

| Table format:<br>Grouped |  | Group A |  |  |  |  |  | Group B |  |  |  |  |  |
| --- | --- | --- | --- | --- | --- | --- | --- | --- | --- | --- | --- | --- | --- |
|  |  | Porites astreoides |  |  |  |  |  | Orbicella franksi |  |  |  |  |  |
|  |  | A:Y1 | A:Y2 | A:Y3 | A:Y4 | A:Y5 | A:Y6 | B:Y1 | B:Y2 | B:Y3 | B:Y4 | B:Y5 | B:Y6 |
| 1 | Control 1 | 477.9956 | 498.9888 | 672.1344 | 768.1330 | 566.9085 | 661.3489 | 634.8769 | 771.4858 | 638.1058 | 435.9589 | 416.4330 | 492.1089 |
| 2 | Control 2 | 485.5405 | 611.5740 | 662.0443 | 625.7935 | 546.8708 | 551.7698 | 549.6299 | 786.8588 | 639.4595 | 442.8147 | 446.1947 | 449.3881 |

Data associated with Figure 3A.

| Table format:<br>Grouped |  | Group A |  |  |  |  |  | Group B |  |  |  |  |  |
| --- | --- | --- | --- | --- | --- | --- | --- | --- | --- | --- | --- | --- | --- |
|  |  | Porites |  |  |  |  |  | Orbicella |  |  |  |  |  |
|  |  | A:Y1 | A:Y2 | A:Y3 | A:Y4 | A:Y5 | A:Y6 | B:Y1 | B:Y2 | B:Y3 | B:Y4 | B:Y5 | B:Y6 |
| 1 | Control | 694.6339 | 486.5243 | 720.1543 | 727.4280 | 576.1440 | 705.1361 | 497.7878 | 567.3501 | 699.9669 | 648.8765 | 492.5736 | 594.3254 |
| 2 | CO<sub>2</sub> | 800.6859 | 639.7489 | 843.2892 | 880.9901 | 635.1358 | 943.2026 | 649.8629 | 629.0223 | 729.7656 | 695.6893 | 579.0927 | 618.0840 |

Data associated with Figure 3B.

| Table format:<br>Grouped |  | Group A |  |  |  |  |  | Group B |  |  |  |  |  |
| --- | --- | --- | --- | --- | --- | --- | --- | --- | --- | --- | --- | --- | --- |
|  |  | Orbicella franksi |  |  |  |  |  | Porites astreoides |  |  |  |  |  |
|  |  | A:Y1 | A:Y2 | A:Y3 | A:Y4 | A:Y5 | A:Y6 | B:Y1 | B:Y2 | B:Y3 | B:Y4 | B:Y5 | B:Y6 |
| 1 | Control | 649.4648 | 773.0557 | 630.4908 | 677.3118 | 635.6335 | 580.4324 | 566.9524 | 664.1805 | 524.4234 | 638.8657 | 653.5860 | 712.0454 |
| 2 | CO<sub>2</sub> | 848.3329 | 923.8812 | 845.9573 | 796.5283 | 848.5911 | 521.5258 | 571.3656 | 885.0912 | 854.0541 | 720.5936 | 677.9351 | 786.4417 |

Data associated with Figure 3C.

Data associated with Figure 3C. *Orbicella franksi*
| Table format:<br>Grouped |  | Group A |  |  |  |  |  |  |  |  |  |  |  |
| --- | --- | --- | --- | --- | --- | --- | --- | --- | --- | --- | --- | --- | --- |
|  |  | Control |  |  |  |  |  |  |  |  |  |  |  |
|  |  | A:Y1 | A:Y2 | A:Y3 | A:Y4 | A:Y5 | A:Y6 | A:Y7 | A:Y8 | A:Y9 | A:Y10 | A:Y11 | A:Y12 |
| 1 | Day 0 | 70.6570 | 50.0640 | 60.6090 | 66.3660 | 64.5390 | 63.9010 | 56.8100 | 66.0330 | 66.3170 | 63.7910 | 62.1820 | 70.8320 |
| 2 | Day 11 | 68.0650 | 58.6300 | 67.2770 | 61.1890 | 58.2310 | 61.3670 | 53.8840 | 58.3750 | 57.9030 | 58.9200 | 57.8460 | 64.8450 |
| Table format:<br>Grouped |  | Group B |  |  |  |  |  |  |  |  |  |  |  |
|  |  | OA |  |  |  |  |  |  |  |  |  |  |  |
|  |  | B:Y2 | B:Y3 | B:Y4 | B:Y5 | B:Y6 | B:Y7 | B:Y8 | B:Y9 | B:Y10 | B:Y11 | B:Y12 |  |
| 1 | Day 0 | 68.9710 | 71.7360 | 67.6840 | 58.8940 | 62.0930 | 61.7680 | 56.7110 | 65.8020 | 67.8980 | 71.1800 | 66.1760 |  |
| 2 | Day 11 | 63.0780 | 67.1910 | 61.9750 | 63.3250 | 62.8020 | 61.1760 | 56.4280 | 60.1100 | 62.6110 | 61.9310 | 62.8830 |  |

*Porites astreoides*
| Table format:<br>Grouped |  | Group A |  |  |  |  |  |  |  |  |  |  |  |
| --- | --- | --- | --- | --- | --- | --- | --- | --- | --- | --- | --- | --- | --- |
|  |  | Control |  |  |  |  |  |  |  |  |  |  |  |
|  |  | A:Y1 | A:Y2 | A:Y3 | A:Y4 | A:Y5 | A:Y6 | A:Y7 | A:Y8 | A:Y9 | A:Y10 | A:Y11 | A:Y12 |
| 1 | Day 0 | 53.5420 | 61.0500 | 59.4890 | 62.2340 | 52.9380 | 49.9050 | 55.4160 | 57.5750 | 57.9120 | 60.0570 | 54.1170 | 58.5200 |
| 2 | Day 11 | 60.9110 | 59.6960 | 56.8780 | 61.2220 | 58.2190 | 48.7170 | 50.1090 | 53.5630 | 59.5640 | 55.1690 | 53.2040 | 53.0340 |
| Table format:<br>Grouped |  | Group B |  |  |  |  |  |  |  |  |  |  |  |
|  |  | OA |  |  |  |  |  |  |  |  |  |  |  |
|  |  | B:Y1 | B:Y2 | B:Y3 | B:Y4 | B:Y5 | B:Y6 | B:Y7 | B:Y8 | B:Y9 | B:Y10 | B:Y11 |  |
| 1 | Day 0 | 58.5600 | 57.3480 | 59.7320 | 59.1700 | 57.2730 | 58.1930 | 53.6020 | 54.3870 | 55.8000 | 59.9480 | 64.5420 |  |
| 2 | Day 11 | 66.5020 | 62.5760 | 59.5230 | 59.8650 | 60.2240 | 53.9760 | 48.6000 | 51.4930 | 53.5640 | 55.7760 | 58.8900 |  |

Data associated with Figure 4.

**Data associated with Figure 4.** Superoxide Dismutase (SOD)
| Table format:<br>Grouped |  | Group A |  |  |  |  |  |  |  |  |  |  |  |
| --- | --- | --- | --- | --- | --- | --- | --- | --- | --- | --- | --- | --- | --- |
|  |  | Orbicella |  |  |  |  |  |  |  |  |  |  |  |
|  |  | A:Y1 | A:Y2 | A:Y3 | A:Y4 | A:Y5 | A:Y6 | A:Y7 | A:Y8 | A:Y9 | A:Y10 | A:Y11 | A:Y12 |
| 1 | Control | 25844.080000 | 10913.600000 | 26255.950000 | 8254.750000 | 27767.440000 |  | 8195.030000 | 9986.120000 | 23622.430000 | 11244.850000 | 38258.330000 | 43782.680000 |
| 2 | CO<sub>2</sub> | 7611.270000 | 11383.570000 | 23431.280000 | 10196.890000 | 24374.170000 | 12004.640000 | 20275.870000 | 35723.370000 | 42177.010000 | 45126.270000 | 32235.770000 | 15069.320000 |

| Table format:<br>Grouped |  | Group B |  |  |  |  |  |  |  |  |  |  |  |
| --- | --- | --- | --- | --- | --- | --- | --- | --- | --- | --- | --- | --- | --- |
|  |  | Porites |  |  |  |  |  |  |  |  |  |  |  |
|  |  | B:Y1 | B:Y2 | B:Y3 | B:Y4 | B:Y5 | B:Y6 | B:Y7 | B:Y8 | B:Y9 | B:Y10 | B:Y11 | B:Y12 |
| 1 | Control | 49647.210000 | 29187.370000 | 67298.830000 | 26686.950000 | 31483.370000 | 32656.390000 | 30894.980000 | 61264.000000 | 18736.590000 | 36258.160000 | 22743.070000 |  |
| 2 | CO<sub>2</sub> | 29464.320000 | 33528.690000 | 21001.980000 | 17781.080000 | 29467.320000 | 29304.130000 | 86622.090000 | 31828.420000 | 34791.930000 | 97514.170000 |  |  |

Catalase (CAT)
| Table format:<br>Grouped |  | Group A |  |  |  |  |  |  |  |  |  |  |  |
| --- | --- | --- | --- | --- | --- | --- | --- | --- | --- | --- | --- | --- | --- |
|  |  | Orbicella |  |  |  |  |  |  |  |  |  |  |  |
|  |  | A:Y1 | A:Y2 | A:Y3 | A:Y4 | A:Y5 | A:Y6 | A:Y7 | A:Y8 | A:Y9 | A:Y10 | A:Y11 | A:Y12 |
| 1 | Control | 0.160000 | 0.110000 | 0.110000 | 0.050000 | 0.180000 |  | 0.050000 | 0.040000 | 0.120000 | 0.070000 | 0.300000 | 0.370000 |
| 2 | CO<sub>2</sub> | 0.100000 | 0.120000 | 0.170000 | 0.120000 | 0.210000 | 0.100000 | 0.350000 | 0.190000 | 0.310000 | 0.300000 | 0.240000 | 0.060000 |
| Table format:<br>Grouped |  | Group B |  |  |  |  |  |  |  |  |  |  |  |
|  |  | Porites |  |  |  |  |  |  |  |  |  |  |  |
|  |  | B:Y1 | B:Y2 | B:Y3 | B:Y4 | B:Y5 | B:Y6 | B:Y7 | B:Y8 | B:Y9 | B:Y10 | B:Y11 | B:Y12 |
| 1 | Control | 0.110000 | 0.090000 | 0.070000 | 0.030000 | 0.020000 | 0.030000 | 0.020000 | 0.180000 | 0.060000 | 0.050000 | 0.040000 |  |
| 2 | CO<sub>2</sub> | 0.010000 | 0.040000 | 0.010000 | 0.010000 | 0.010000 | 0.080000 | 0.140000 | 0.040000 | 0.100000 | 0.110000 |  |  |

Reduced Glutathione (GSH)
| Table format:<br>Grouped |  | Group A |  |  |  |  |  |  |  |  |  |  |  |
| --- | --- | --- | --- | --- | --- | --- | --- | --- | --- | --- | --- | --- | --- |
|  |  | Orbicella |  |  |  |  |  |  |  |  |  |  |  |
|  |  | A:Y1 | A:Y2 | A:Y3 | A:Y4 | A:Y5 | A:Y6 | A:Y7 | A:Y8 | A:Y9 | A:Y10 | A:Y11 | A:Y12 |
| 1 | Control | 33.626760 | 13.36912 | 24.556450 | 15.27778 | 38.858700 | 9.840838 | 9.980769 | 51.695740 | 7.448083 | 55.049670 | 98.19915 |  |
| 2 | CO<sub>2</sub> | 9.173820 | 18.10233 | 35.500000 | 19.49850 | 39.293980 | 26.171880 | 37.816460 | 53.839870 | 55.729170 | 112.771700 | 66.80463 | 22.80000 |
| Table format:<br>Grouped |  | Group B |  |  |  |  |  |  |  |  |  |  |  |
|  |  | Porites |  |  |  |  |  |  |  |  |  |  |  |
|  |  | B:Y1 | B:Y2 | B:Y3 | B:Y4 | B:Y5 | B:Y6 | B:Y7 | B:Y8 | B:Y9 | B:Y10 | B:Y11 | B:Y12 |
| 1 | Control | 52.0000 | 41.3000 | 70.0500 | 37.0600 | 35.6000 | 28.9800 | 43.3500 | 98.9800 | 24.3500 | 32.9500 | 28.6800 |  |
| 2 | CO<sub>2</sub> | 29.7200 | 21.5000 | 19.8800 | 21.5100 | 24.3900 | 23.4600 | 116.9600 | 47.8900 | 36.2300 | 102.2900 |  |  |

Glutathione S-Transferase (GST)
| Table format:<br>Grouped |  | Group A |  |  |  |  |  |  |  |  |  |  |  |
| --- | --- | --- | --- | --- | --- | --- | --- | --- | --- | --- | --- | --- | --- |
|  |  | Orbicella |  |  |  |  |  |  |  |  |  |  |  |
|  |  | A:Y1 | A:Y2 | A:Y3 | A:Y4 | A:Y5 | A:Y6 | A:Y7 | A:Y8 | A:Y9 | A:Y10 | A:Y11 | A:Y12 |
| 1 | Control | 0.450000 | 0.17000 | 0.350000 | 0.30000 | 0.510000 |  | 0.150000 | 0.210000 | 0.360000 | 0.240000 | 0.61000 | 0.61000 |
| 2 | CO<sub>2</sub> | 0.170000 | 0.32000 | 0.760000 | 0.63000 | 0.680000 | 0.480000 | 0.380000 | 0.560000 | 0.760000 | 0.230000 | 0.15000 | 0.06000 |
| Table format:<br>Grouped |  | Group B |  |  |  |  |  |  |  |  |  |  |  |
|  |  | Porites |  |  |  |  |  |  |  |  |  |  |  |
|  |  | B:Y1 | B:Y2 | B:Y3 | B:Y4 | B:Y5 | B:Y6 | B:Y7 | B:Y8 | B:Y9 | B:Y10 | B:Y11 | B:Y12 |
| 1 | Control | 0.3200 | 0.2200 | 0.2400 | 0.1000 | 0.1900 | 0.1100 | 0.1700 | 0.5800 | 0.0600 | 0.1100 | 0.1100 |  |
| 2 | CO<sub>2</sub> | 0.1300 | 0.0700 | 0.0700 | 0.0400 | 0.1300 | 0.1100 | 0.2500 | 0.1600 | 0.1700 | 0.4500 |  |  |

TBARS (lipid peroxidation)
| Table format:<br>Grouped |  | Group A |  |  |  |  |  |  |  |  |  |  |  |
| --- | --- | --- | --- | --- | --- | --- | --- | --- | --- | --- | --- | --- | --- |
|  |  | Orbicella franksi |  |  |  |  |  |  |  |  |  |  |  |
|  |  | A:Y1 | A:Y2 | A:Y3 | A:Y4 | A:Y5 | A:Y6 | A:Y7 | A:Y8 | A:Y9 | A:Y10 | A:Y11 | A:Y12 |
| 1 | Control | 41.278440 | 25.33035 | 34.708440 | 16.51456 | 39.406350 | 16.017680 | 13.721890 | 46.533990 | 18.819120 | 64.760570 | 81.06258 |  |
| 2 | CO<sub>2</sub> | 17.010560 | 29.96712 | 48.461540 | 28.62160 | 40.331200 | 38.902240 | 51.752680 | 72.360480 | 81.442310 | 68.770900 | 68.96332 | 26.55000 |
| Table format:<br>Grouped |  | Group B |  |  |  |  |  |  |  |  |  |  |  |
|  |  | Porites astreoides |  |  |  |  |  |  |  |  |  |  |  |
|  |  | B:Y1 | B:Y2 | B:Y3 | B:Y4 | B:Y5 | B:Y6 | B:Y7 | B:Y8 | B:Y9 | B:Y10 | B:Y11 | B:Y12 |
| 1 | Control | 38.6400 | 31.9500 | 38.7700 | 27.7700 | 53.1100 | 30.2800 | 37.8100 | 50.1100 | 19.4700 | 34.0200 | 29.1100 |  |
| 2 | CO<sub>2</sub> | 23.1000 | 24.0600 | 21.3300 | 20.1100 | 23.8400 | 17.6600 | 47.9200 | 28.2100 | 31.0400 | 58.4700 |  |  |

Carbonyl (protein oxidation)
| Table format:<br>Grouped |  | Group A |  |  |  |  |  |  |  |  |  |  |  |
| --- | --- | --- | --- | --- | --- | --- | --- | --- | --- | --- | --- | --- | --- |
|  |  | Orbicella |  |  |  |  |  |  |  |  |  |  |  |
|  |  | A:Y1 | A:Y2 | A:Y3 | A:Y4 | A:Y5 | A:Y6 | A:Y7 | A:Y8 | A:Y9 | A:Y10 | A:Y11 | A:Y12 |
| 1 | Control | 24.701240 | 28.83900 | 37.646630 | 13.99589 | 21.245060 | 13.534020 | 10.541960 | 46.511630 | 10.492300 | 50.421430 | 33.51310 |  |
| 2 | CO<sub>2</sub> | 15.184030 | 19.87164 | 40.959590 | 21.06015 | 31.250000 | 26.199500 | 35.193710 | 53.847300 | 59.375000 | 48.089590 | 54.93679 |  |

| Table format:<br>Grouped |  | Group B |  |  |  |  |  |  |  |  |  |  |  |
| --- | --- | --- | --- | --- | --- | --- | --- | --- | --- | --- | --- | --- | --- |
|  |  | Porites |  |  |  |  |  |  |  |  |  |  |  |
|  |  | B:Y1 | B:Y2 | B:Y3 | B:Y4 | B:Y5 | B:Y6 | B:Y7 | B:Y8 | B:Y9 | B:Y10 | B:Y11 | B:Y12 |
| 1 | Control | 140.08000 | 75.29736 | 90.82734 | 37.72727 | 135.63950 | 95.37515 | 88.58749 | 71.35307 | 88.18182 | 54.50000 | 88.57189 |  |
| 2 | CO<sub>2</sub> | 20.31302 | 29.03311 | 28.63162 | 17.82418 | 28.08748 | 17.11076 | 57.62987 | 41.22749 | 33.28334 | 72.61641 | 11.08000 |  |

Data associated with Figure 5.

